# *AmoA*-targeted polymerase chain reaction primers for the specific detection and quantification of comammox *Nitrospira* in the environment

**DOI:** 10.1101/096891

**Authors:** Petra Pjevac, Clemens Schauberger, Lianna Poghosyan, Craig W. Herbold, Maartje A. H. J. van Kessel, Anne Daebeler, Michaela Steinberger, Mike S. M. Jetten, Sebastian Lücker, Michael Wagner, Holger Daims

## Abstract

Nitrification, the oxidation of ammonia via nitrite to nitrate, has always been considered to be catalyzed by the concerted activity of ammonia and nitrite-oxidizing microorganisms. Only recently, complete ammonia oxidizers (‘comammox’), which oxidize ammonia to nitrate on their own, were identified in the bacterial genus *Nitrospira*, previously assumed to contain only canonical nitrite oxidizers. *Nitrospira* are widespread in nature, but for assessments of the distribution and functional importance of comammox *Nitrospira* in ecosystems, cultivation-independent tools to distinguish comammox from strictly nitrite-oxidizing *Nitrospira* are required. Here we developed new PCR primer sets that specifically target the *amoA* genes coding for subunit A of the distinct ammonia monooxygenase of comammox *Nitrospira*. While existing primers capture only a fraction of the known comammox *amoA* diversity, the new primer sets cover as much as 95% of the comammox *amoA* clade A and 92% of the clade B sequences in a reference database containing 326 comammox *amoA* genes with sequence information at the primer binding sites. Application of the primers to 13 samples from engineered systems (a groundwater well, drinking water treatment and wastewater treatment plants) and other habitats (rice paddy and forest soils, rice rhizosphere, brackish lake sediment and freshwater biofilm) detected comammox *Nitrospira* in all samples and revealed a considerable diversity of comammox in most habitats. Excellent primer specificity for comammox *amoA* was achieved by avoiding the use of highly degenerate primer preparations and by using equimolar mixtures of oligonucleotides that match existing comammox *amoA* genes. Quantitative PCR with these equimolar primer mixtures was highly sensitive and specific, and enabled the efficient quantification of clade A and clade B comammox *amoA* gene copy numbers in environmental samples. The measured relative abundances of comammox *Nitrospira*, compared to canonical ammonia oxidizers, were highly variable across environments. The new comammox *amoA*-targeted primers will enable more encompassing studies of nitrifying microorganisms in diverse ecosystems.

## 1 Introduction

Nitrification is an essential process of the global biogeochemical nitrogen cycle and plays a pivotal role in biological wastewater treatment and in drinking water production. The recent discovery of the first complete ammonia oxidizers (‘comammox’) in the bacterial genus *Nitrospira* (Daims *et al.*, 2015; van Kessel *et al.*, 2015) was highly unexpected since *Nitrospira* were always regarded as canonical nitrite-oxidizing bacteria (NOB) (Watson *et al.*, 1986; Ehrich *et al.*, 1995; Spieck *et al.*, 2006; Lebedeva *et al.*, 2008; 2011). This discovery has raised a number of important questions, such as how often comammox *Nitrospira* occur in nitrifying microbial communities, and how relevant they are for nitrification compared to ammonia-oxidizing bacteria (AOB), archaea (AOA), and NOB.

Members of the genus *Nitrospira* have been assigned to at least six sublineages (Daims *et al.*, 2001; Lebedeva *et al.*, 2008; 2011). They are widespread in virtually all oxic natural and engineered ecosystems, and an impressively high diversity of coexisting uncultured *Nitrospira* strains has been detected in wastewater treatment plants and soils (Pester *et al.*,2014; Gruber-Dorninger *et al.*, 2015). All known comammox organisms belong to *Nitrospira* sublineage II (Daims *et al.*, 2015; van Kessel *et al.*, 2015; Pinto *et al.*, 2015; Palomo *et al.*,2016). This sublineage contains also canonical NOB, which lack the genes for ammonia oxidation (Daims *et al.*, 2001, Koch *et al.*, 2015). Moreover, the known comammox *Nitrospira* do not form a monophyletic clade within *Nitrospira* lineage II in phylogenies based on 16S rRNA genes or *nxrB*, the gene encoding subunit beta of the functional key enzyme nitrite oxidoreductase. Instead, they intersperse with the strict NOB in these phylogenetic trees (Daims *et al.*, 2015; van Kessel *et al.*, 2015; Pinto *et al.*, 2015). Finally, it remains unknown whether other *Nitrospira* sublineages (Daims *et al.*, 2001; Lebedeva *et al.*,2008, 2011) also contain comammox members. Consequently, it is impossible to infer from 16S rRNA or *nxrB* gene phylogenies whether yet uncharacterized *Nitrospira* bacteria are comammox or strict NOB, although such attempts have been published (Gonzalez-Martinez *et al.*, 2016).

Intriguingly, comammox *Nitrospira* possess novel types of ammonia monooxygenase (AMO) and hydroxylamine oxidoreductase (HAO), the key enzymes of aerobic ammonia oxidation. The comammox AMO is phylogenetically distinct from the AMO forms of canonical AOB and AOA (Daims *et al.*, 2015; van Kessel *et al.*, 2015). The *amoA* gene encoding AMO subunit A has become a widely used functional and phylogenetic marker gene for bacterial and archaeal ammonia oxidizers (Rotthauwe *et al.*, 1997; Purkhold *et al.*, 2000; Juniper *et al.*,2008; Pester *et al.*, 2012). Public database mining for sequences related to the unique comammox *amoA* revealed the presence of putative comammox organisms in various environments including soils (paddy rice soils, other agricultural soils, forest soils, grassland soils), freshwater habitats (wetlands, rivers, lakes, groundwater basins), groundwater wells (GGWs), full-scale wastewater treatment plants (WWTPs), and drinking water treatment plants (DWTPs) (Daims *et al.*, 2015; van Kessel *et al.*, 2015). While this provides strong indications of a broad habitat range of comammox organisms, current knowledge about the environmental distribution and abundance of comammox *Nitrospira* is very limited and needs to be explored.

The *amoA* genes of comammox *Nitrospira* form two monophyletic sister clades (Daims *et al.*, 2015; van Kessel *et al.*, 2015), which are referred to as clade A and clade B. Clade A also contains some genes that were previously assigned to the methanotroph *Crenothrix polyspora* (Stoecker *et al.*, 2006), but this assignment has recently been corrected (Oswald *et al.*, in revision). Established PCR primer sets specifically targeting the *amoA* genes of AOB or AOA (Sinigalliano *et al.*, 1995; Rotthauwe *et al.*, 1997; Juretschko *et al.*, 1998; Stephen *et al.*, 1999; Nold *et al.*, 2000; Norton et al., 2002; Webster *et al.*, 2002; Okano *et al.*, 2004; Francis et al., 2005; Treusch *et al.*, 2005; Mincer *et al.*, 2007; Juniper *et al.*, 2008; Meinhardt *et al.*, 2015) do not amplify any clade A and clade B comammox *amoA.* Co-amplification of some comammox *amoA* genes occurs with a primer set targeting both betaproteobacterial *amoA* and the A subunit of the particulate methane monooxygenase *(pmoA* of pMMO; Holmes *et al.*, 1995; Luesken *et al.*, 2011). These primers, however, only target a fraction of the known comammox *amoA* genes. A recently established two-step PCR protocol (Wang *et al.*, 2016) relies on the forward primer published by Holmes and colleagues (1995) and a new, highly degenerate reverse primer. It detects the bacterial copper-containing monooxygenase (CuMMO) genes including *pmoA*, betaproteobacterial *amoA*, and at least the comammox *amoA* genes that are amplified by the Holmes forward primer. However, because of its broad coverage of the CuMMOs, this primer pair does not allow a specific detection or quantification by quantitative PCR (qPCR) of comammox *amoA* genes. The two sets of comammox *amoA* targeted (q)PCR primers recently published by Bartelme and colleagues (2017) are, in contrast, highly specific. These primers only detect a small fraction of available comammox *amoA* gene sequences within clade A, and cannot be used to amplify comammox clade B *amoA* genes.

To enable the direct detection and quantification of comammox *amoA* genes in environmental samples, we designed in this study two new *amoA*-targeted primer sets specific for clade A or clade B comammox *amoA* genes, respectively. Subsequently, we applied these new primers to efficiently screen various habitats for the presence of comammox organisms and to rapidly quantify the abundances of comammox in selected samples.

## 2 Materials and Methods

### 2.1 Database mining and sequence collection

The amino acid sequences of bacterial AmoA and PmoA were extracted from publicly available metagenomic datasets stored in the Integrated Microbial Genomes databases (IMGER and -MER) using the functional profiler tool against a specific bacterial AmoA/PmoA pfam (PF02461). For characterization of comammox AmoA, betaproteobacterial AmoA, and PmoA, sequences collected from the Integrated Microbial Genomes databases were augmented with nearly full length amino acid sequences collected from the Pfam site and from NCBI Genbank as described in Daims *et al.* (2015). Collected sequences were filtered against a hidden Markov model (hmm) using hmmsearch (http://hmmer.janelia.org/) with the AmoA/PmoA hmm (PF02461) requiring an expect value of <0.0001, and clustered at 90% identity using USEARCH (Edgar, 2010). Cluster centroids were aligned using Mafft (Katoh *et al.*, 2002) and phylogenetic affiliation with comammox AmoA or betaproteobacterial AmoA was verified with phylogenetic trees calculated using FastTree (Prince *et al.*, 2009). Cluster centroids were expanded and reclustered at 99% identity with USEARCH and the phylogenetic affiliation with comammox and/or betaproteobacterial AmoA was re-assessed using FastTree.

### 2.2 Phylogenetic analyses

For amino acid sequences that clustered with comammox AmoA and a selection of outgroup sequences affiliating with bacterial AmoA and PmoA, the corresponding nucleic acid sequences were recovered and aligned according to their amino acid translations using MUSCLE (Edgar, 2004). These nucleotide sequence alignments were manually corrected for frameshifts and used for calculating maximum likelihood trees with RaxmlHPC (Stamatakis *et al.*, 2005) using the GTRCAT approximation of rate heterogeneity and 1,000 bootstrap iterations. The generated reference tree was imported into the ARB software package (Ludwig *et al.*, 2004) and used as reference tree for the subsequent addition of short and nearly full length comammox *amoA*, betaproteobacterial *amoA* and *pmoA* nucleotide sequences that were recovered from public databases. These sequences were added to the reference tree without changes of the overall tree topology by using the “parsimony interactive” tool of ARB.

### 2.3 Primer design and PCR

Two degenerate PCR primer pairs (Table 1) targeting clade A or clade B comammox *amoA* genes, respectively, were designed and evaluated using the *amoA/pmoA* reference sequence dataset and the ‘probe design’ and ‘probe match’ functions of ARB (Ludwig *et al.*, 2004). The oligonucleotide primers were obtained from Biomers (Ulm, Germany).

**Table 1.**
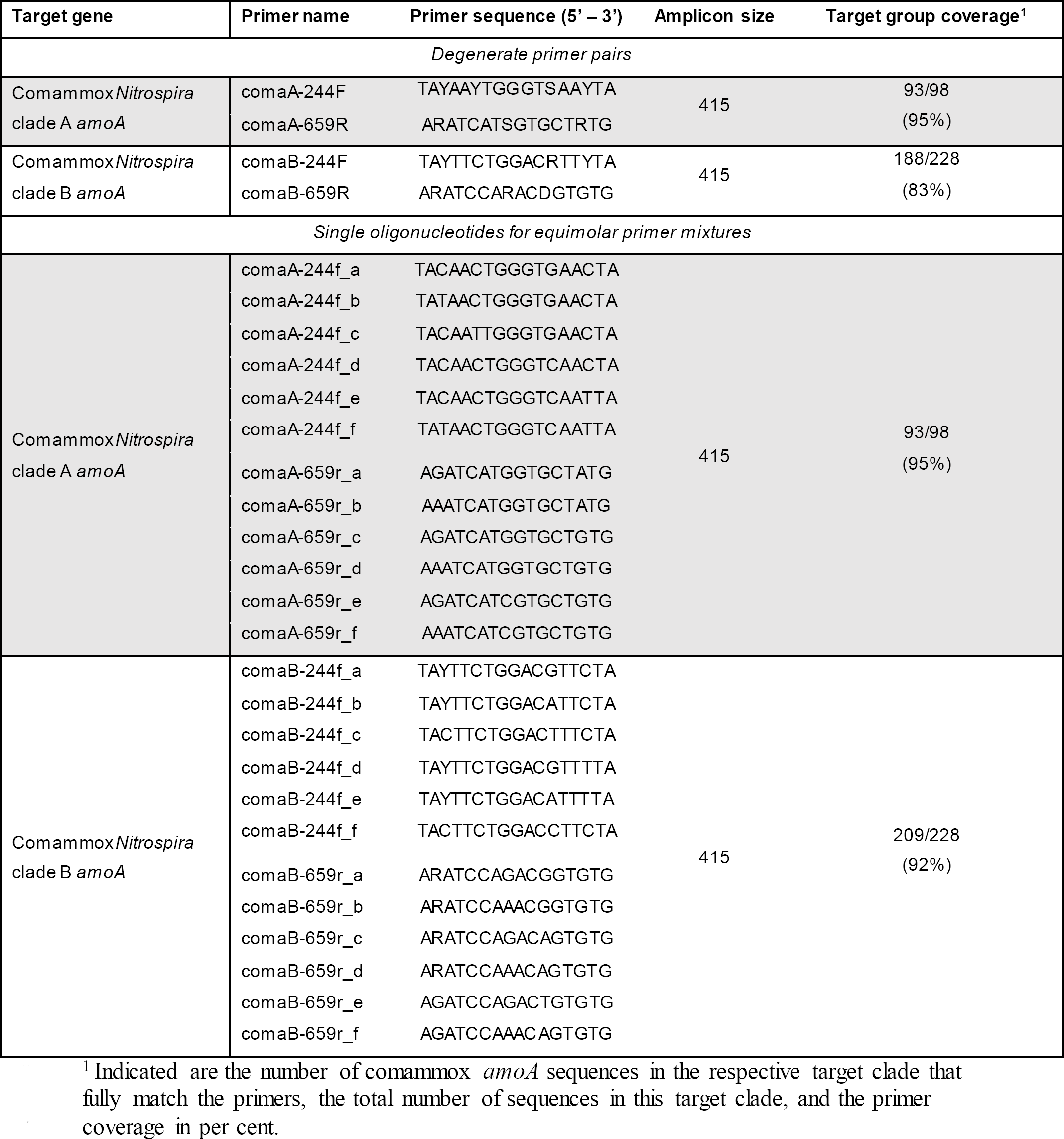
Comammox *amoA*-targeted primers designed in this study, and in silico evaluation of the primer coverage of the target clades in our comammox amoA reference sequence databas

The optimal annealing temperature for the primers was determined by temperature gradient PCR using total genomic DNA extracted by a phenol-chloroform based method (details below) from a GWW sample (Wolfenbüttel, Germany), which contained both clade A and clade B comammox *Nitrospira* (Daims *et al.*, 2015). An annealing temperature range from 42 to 52 °C was chosen for experimental evaluation based on the theoretical melting temperature of 48 °C of the designed primers (http://biotools.nubic.northwestern.edu/OligoCalc.html). Thermal cycling was carried out with an initial denaturation step at 94 °C for 5 min, which was followed by 25 cycles of denaturation at 94 °C for 30 s, primer annealing at 42 to 52 °C for 45 s, and elongation at 72 °C for 1 min. Cycling was completed by a final elongation step at 72 °C for 10 min. The optimal annealing temperature for both primer pairs was found to be 52 °C. At lower annealing temperatures, unspecific amplification of DNA fragments shorter than the expected amplicon length (415 bp) was observed. Since the amplification efficiency of correctly sized amplicons was lower at 52 °C than at 48–50 °C, temperatures above 52 °C were not evaluated. The PCR reactions were performed in a DreamTaq Green PCR mix with 1× Dream Taq Green Buffer containing 2 mM MgCl_2_, 0.025 U DreamTaq DNA polymerase, 0.2 mM dNTPs, 0.5 μM primers and 0.1 mg/mL bovine serum albumin (Fermentas, Thermo Fischer Scientific, Waltham, MA, USA).

### 2.4 Screening of environmental samples for comammox amoA

Total nucleic acids were extracted from the environmental samples (Table 2) by using i) the Fast DNA SPIN Kit for soil (MP Biomedicals, Santa Ana, CA, USA), ii) the PowerSoil DNA Isolation Kit (MO BIO Laboratories, Carlsbad, CA, USA), or iii) a phenol-chloroform based total nucleic acid extraction protocol described by Angel *et al.*, (2012). PCRs were performed with either the DreamTaq Green PCR mix as described above, or with the PerfeCTa SYBR green FastMix PCR premix containing 1.5 mM of MgCl_2_ (Quanta Bioscience, Beverly, MA, USA) and 0.1 μM primers. Amplicons of the expected length (415 bp) were purified by using the QIAquick PCR purification Kit (Qiagen, Hilden, Germany). Alternatively, the QIAquick Gel extraction Kit (Qiagen, Hilden, Germany) was used to purify PCR products of the correct length from agarose gels if additional, unspecific amplification products of different lengths were present. The purified PCR products were cloned in *E. coli* with the TOPO-TA cloning kit (Invitrogen, Karlsruhe, Germany) or the pGEM-T Easy Vector System (Promega, Mannheim, Germany) by following the manufacturer's instructions. PCR products obtained from cloned vector inserts from the TOPO-TA kit were Sanger-sequenced with the M13F primer at Microsynth (Balgach, Switzerland). For clones generated with the pGEM-TEasy Vector SysteM, plasmids were extracted with the GeneJET plasmid miniprep kit (Thermo Fischer Scientific, Waltham, MA, USA) and the inserts were Sanger-sequenced with the M13F primer at Baseclear (Leiden, the Netherlands). The recovered sequences were quality checked and vector trimmed using the Sequencher version 4.6.1 (GeneCodes Corporation, Ann Arbor, MI, USA) or the Chromas version 2.6.1 (Technelysium Pty Ltd, Brisbane, Australia) software packages, and were added as described above to the *amoA/pmoA* nucleotide sequence reference tree in ARB.

**Table 2.**
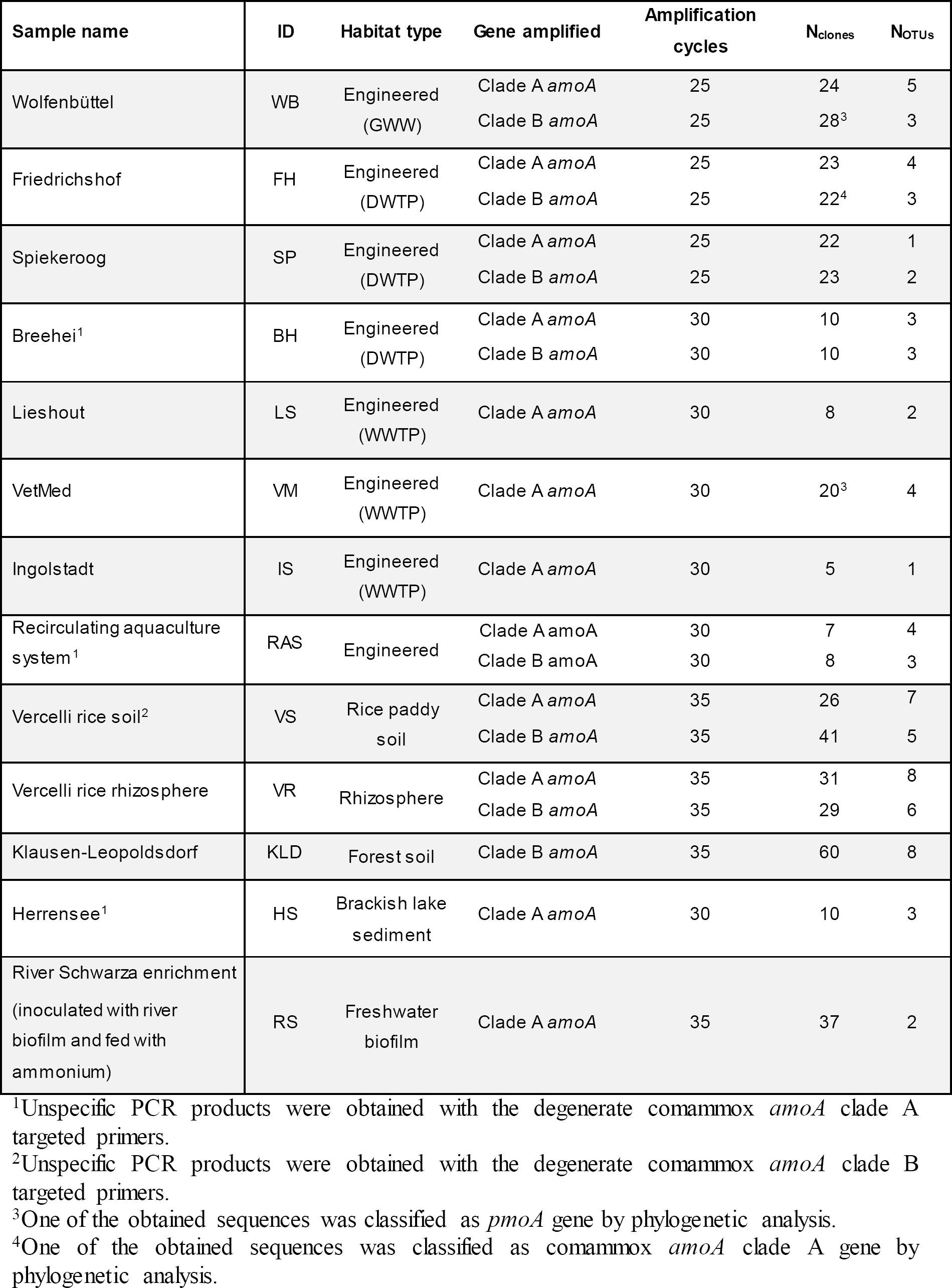
Environmental samples analyzed in this study, PCR amplification cycles, and numbers of cloned comammox amoA sequences and OTUs retrieved from each sample.

| Sample name | ID | Habitat type | Gene amplified | Amplification cycles | N<sub>clones</sub> | N<sub>OTUs</sub> |
| --- | --- | --- | --- | --- | --- | --- |
| Wolfenbüttel | WB | Engineered (GWW) | Clade A *amoA* | 25 | 24 | 5 |
|  |  |  | Clade B *amoA* | 25 | 28<sup>3</sup> | 3 |
| Friedrichshof | FH | Engineered (DWTP) | Clade A *amoA* | 25 | 23 | 4 |
|  |  |  | Clade B *amoA* | 25 | 22<sup>4</sup> | 3 |
| Spiekeroog | SP | Engineered (DWTP) | Clade A *amoA* | 25 | 22 | 1 |
|  |  |  | Clade B *amoA* | 25 | 23 | 2 |
| Breehei<sup>1</sup> | BH | Engineered (DWTP) | Clade A *amoA* | 30 | 10 | 3 |
|  |  |  | Clade B *amoA* | 30 | 10 | 3 |
| Lieshout | LS | Engineered (WWTP) | Clade A *amoA* | 30 | 8 | 2 |
| VetMed | VM | Engineered (WWTP) | Clade A *amoA* | 30 | 20<sup>3</sup> | 4 |
| Ingolstadt | IS | Engineered (WWTP) | Clade A *amoA* | 30 | 5 | 1 |
| Recirculating aquaculture system<sup>1</sup> | RAS | Engineered | Clade A *amoA* | 30 | 7 | 4 |
|  |  |  | Clade B *amoA* | 30 | 8 | 3 |
| Vercelli rice soil<sup>2</sup> | VS | Rice paddy soil | Clade A *amoA* | 35 | 26 | 7 |
|  |  |  | Clade B *amoA* | 35 | 41 | 5 |
| Vercelli rice rhizosphere | VR | Rhizosphere | Clade A *amoA* | 35 | 31 | 8 |
|  |  |  | Clade B *amoA* | 35 | 29 | 6 |
| Klausen-Leopoldsdorf | KLD | Forest soil | Clade B *amoA* | 35 | 60 | 8 |
| Herrensee<sup>1</sup> | HS | Brackish lake sediment | Clade A *amoA* | 30 | 10 | 3 |
| River Schwarza enrichment (inoculated with river biofilm and fed with ammonium) | RS | Freshwater biofilm | Clade A *amoA* | 35 | 37 | 2 |

<sup>1</sup>Unspecific PCR products were obtained with the degenerate comammox *amoA* clade A targeted primers.
<sup>2</sup>Unspecific PCR products were obtained with the degenerate comammox *amoA* clade B targeted primers.
<sup>3</sup>One of the obtained sequences was classified as *pmoA* gene by phylogenetic analysis.
<sup>4</sup>One of the obtained sequences was classified as comammox *amoA* clade A gene by phylogenetic analysis.

### 2.5 Quantitative PCR

Quantification of comammox *amoA* genes was performed with individually prepared equimolar primer mixes for comammox *amoA* clade A and clade B genes (Table 1) on a selection of samples from different habitats (Table S2). Amplification of comammox *amoA* genes was performed with 3 min initial denaturation at 95 °C, followed by 45 cycles of 30 sec at 95 °C, 45 sec at 52 °C, and 1 min at 72 °C. Fluorescence was measured at 72 °C for amplicon quantification. After amplification, an amplicon melting curve was recorded in 0.5 °C steps between 38 and 96 °C. Amplification of archaeal and betaproteobacterial *amoA* genes was performed with the GenAOA and the RottAOB primers as described by Meinhardt *et al.* (2015). All qPCR assays were performed on a Bio-Rad C1000 CFX96 Real-Time PCR system (Bio-Rad, Hercules, CA, USA) in Bio-Rad iQ SYBR Green Supermix (Bio-Rad, Hercules, CA, USA), containing 50 U/ml iTaq DNA polymerase, 0.4 mM dNTPs, 100 mM KCl, 40 mM Tris-HCl, 6 mM MgCl_2_, 20 mM fluorescein, and stabilizers. The respective equimolar comammox *amoA* primer mixtures (Table 1), or archaeal or bacterial *amoA* primers were added to a final concentration of 0.5 μM. Environmental DNA samples were added to each PCR reaction to a final volume of 20 μl reaction mix. Different template DNA dilutions (10x to 10,000x) resulting in DNA concentrations from 0.004 to 24 ng were applied to minimize possible PCR inhibition caused by excess DNA or co-extracted substances.

All assays were performed in triplicate for each dilution. For each assay, triplicate standard series were generated by tenfold serial dilutions (10^1^-10^8^ gene copies/μl) from purified M13- PCR products obtained from cloned vector inserts generated with the TOPO-TA cloning kit as described above. Clones of comammox *amoA* genes used for standard curve generation originated from this study, while the betaproteobacterial or archaeal *amoA* genes for the respective standard curves were obtained from pure cultures of *Nitrosomonas nitrosa* Nm90 and *Nitrososphera gargensis*, respectively. The correlation coefficient (r^2^) for each of the external standard curves was ≥0.98. The amplification efficiency for comammox *amoA* clade A and clade B genes (standard curves and environmental samples) was 88.5% and 87.0%, respectively. Amplification efficiency for archaeal *amoA* genes was 88.7%, and 92.3% for betaproteobacterial *amoA* genes.

## 3 Results and Discussion

### 3.1 Design and evaluation of comammox amoA-specific PCR primers

A principal goal of this study was to develop a specific amoA-targeted PCR assay for the efficient and cultivation-independent detection and quantification of comammox *Nitrospira* in environmental samples. Mining of public databases retrieved only a small number (376) of sequences related to clade A or clade B comammox *amoA*, and only 32 of these sequences cover almost the full length of the comammox *amoA* genes (840-846 bp for clade A, 885-909 bp for clade B according to the available genomic and metagenomic datasets from comammox *Nitrospira).* To achieve a good coverage of the known comammox *amoA* gene diversity, primer design was restricted to the range of aligned nucleotide positions for which sequence information exists in at least 80% of the publicly available comammox *amoA* sequences. Furthermore, availability of sequence information for at least one of the primer binding sites in the great majority of betaproteobacterial *amoA* gene sequences (14,000/14,019) and bacterial *pmoA* gene sequences (9,913/10,289) facilitated the design of highly comammox amoA-specific primers. Two degenerate primer pairs (ComaA-244F/659R and ComaB-244F/659R) targeted 95% and 83% of the *amoA* gene sequences from comammox clade A or B, respectively (Table 1). *In silico* analyses confirmed that each primer contained at least 4 mismatches to all sequences within the respective other comammox *amoA* clade. Furthermore, at least 3 mismatches to almost all sequences affiliated with betaproteobacterial *amoA* and with *pmoA* genes were present. Merely 23 out of 14,019 betaproteobacterial *amoA* had only two mismatches to the comammox *amoA* clade A targeted primers. Further, only 22 and 2 out of 9,913 *pmoA* gene sequences showed only two mismatches to the comammox *amoA* clade A and clade B-targeted primers, respectively. No betaproteobacterial *amoA* and no *pmoA* gene sequences had less than two mismatches to any of the comammox amoA-targeted primer pairs.

The new primer sets were first evaluated with a pasty iron sludge sample from the GWW Wolfenbüttel, which was known to harbor both clade A and clade B comammox *Nitrospira* according to a previous metagenomic analysis (Daims *et al.*, 2015). A single amplicon of the expected length (415 bp) was obtained from this sample with both primer pairs after PCR with an annealing temperature of 52 °C. Cloning and sequencing of the PCR products exclusively retrieved sequences of comammox *amoA* from clade A or clade B, respectively (Fig. 1). After application of the two primer sets to 12 additional samples from various environments, comammox *amoA* clade A was detected in 11 samples and clade B in 7 samples (see below and Table 2). Single, correctly sized amplicons were obtained from the majority of samples (8 of the clade A-positive samples and 6 of the clade B-positive samples, Table 2). In the remaining cases, unspecific PCR products of different lengths were observed in addition to the expected *amoA* amplicons. These unspecific PCR products appeared irrespectively of the applied DNA extraction method and the number of PCR cycles, but were overcome by either purifying the target *amoA* amplicon by agarose gel excision (for the lake Herrensee sample) or by a PCR-based clone screening for the right insert size after cloning (for the remaining samples with unspecific PCR products). In summary, we retrieved 446 CuMMO sequences in this study by PCR, cloning and sequencing from the 13 environmental samples (Table 2), 444 of which were comammox *amoA* gene sequences. Only two sequences did not cluster with comammox *amoA* clades, but were classified as *pmoA.* Furthermore, one sequence obtained with the comammox *amoA* clade B-targeted primers turned out to be affiliated with comammox *amoA* clade A. Thus, the degenerate comammox amoA-targeted primers offer a straightforward, fast and robust approach for the detection and identification of comammox *Nitrospira* in complex samples. Notably, with an amplicon size of 415 bp the primers are also suitable for high-throughput amplicon sequencing by current Illumina technologies.

**Figure. 1.**
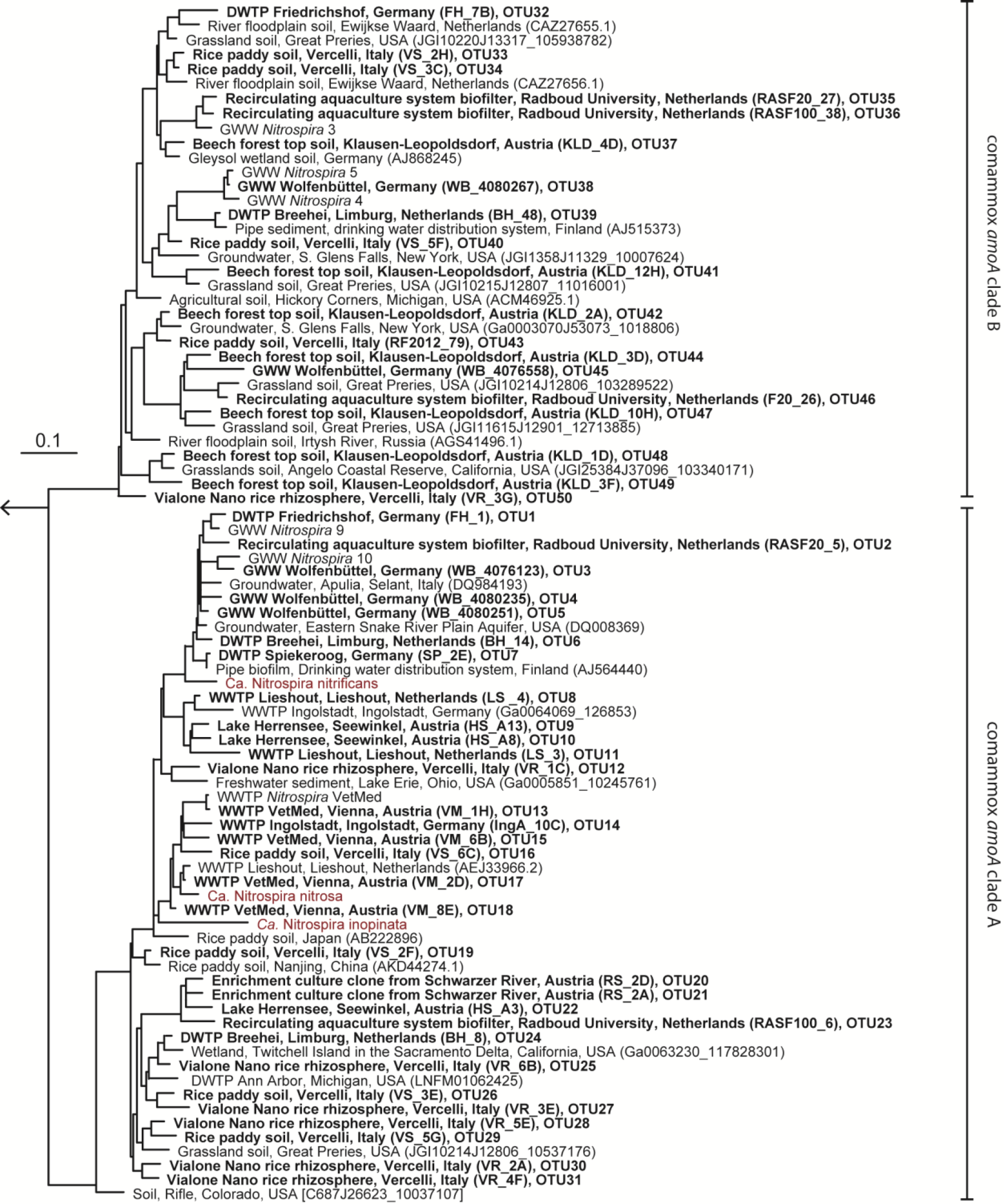
Maximum likelihood tree showing the phylogenetic affiliation of comammox *amoA* genes obtained in this study (printed in boldface) to reference *amoA* sequences from comammox clades A and B. One representative sequence from each OTU is shown. Identifiers of clones (in parentheses) and of OTUs are indicated. Previously described comammox *Nitrospira* strains are shown in red. The outgroup consisted of selected betaproteobacterial amoA and proteobacterial pmoA genes. The scale bar indicates the estimated change rate per nucleotide.

Since amplification of non-target DNA was observed with some samples, we took into account that primer degeneracy may impair PCR specificity (Linhart and Shamir, 2002). Furthermore, primer degeneracy can cause an uneven amplification of different sequence variants (Polz and Cavanaugh, 1998). In general, less degenerate, defined primer mixtures can reduce unspecific primer binding and yield cleaner amplification products from complex environmental samples (Linhart and Shamir, 2002). This is of particular importance for obtaining correct population size estimates in qPCR assays. Consequently, less degenerate primer mixtures consisting of separately synthesized versions of the respective forward and reverse primer, with each primer version containing only one or no base ambiguity (Table 1), were also tested. All selected primer versions in these mixtures matched real comammox *amoA* gene sequences in our database. In contrast, the original (more degenerate) primer sets inevitably contained also co-synthesized oligonucleotide versions that did not match any known target comammox *amoA* sequence. The less degenerate mixtures offered the same coverage of comammox *amoA* clade A (95%), and by adding one additional oligonucleotide to the forward primer mix we increased the coverage of comammox *amoA* clade B from 83 to 92%. Considerably improved PCR results were achieved by applying these primer mixtures to the samples prone to unspecific amplification, as only amplicons of the expected size were obtained after PCR (data now shown). Based on their improved specificity and the higher coverage of comammox *amoA* clade B, use of the defined, manually pooled equimolar primer mixtures (Table 1) is recommended. For comammox amoA-targeted qPCR experiments that use the primer sets presented here, application of the less degenerate, equimolar primer mixtures is mandatory.

### 3.2 Environmental detection and diversity of comammox Nitrospira

Intriguingly, comammox *amoA* genes were detected in all samples (Table 2) encompassing forest soil, rice paddy soils and rice rhizosphere, a freshwater biofilm and brackish lake sediment, as well as WWTPs and DWTPs. After cloning and sequencing, the obtained *amoA* sequences were clustered in operational taxonomic units (OTUs) based on a sequence identity threshold of 95% (Francis *et al.*, 2003). It should be noted that these OTUs might not delineate species, as an appropriate species-level sequence identity cutoff for comammox *amoA* remains unknown. This would need to be determined by correlating *amoA* sequence identities with 16S rRNA identities or with genome-wide average nucleotide identities (Purkhold *et al.*, 2000; Richter and Roselló-Mora, 2009; Pester *et al.*, 2014) once more genomic data from comammox *Nitrospira* become available. A phylogenetic analysis of the *amoA* gene OTU representatives revealed a substantial diversity of comammox *Nitrospira* in almost all samples (Fig. 1, Table 2, Table S1). Both clade A and clade B comammox *Nitrospira* were detected in seven samples. Only clade A members were found in the WWTPs, the brackish lake sediment, and the river biofilm enrichment, whereas we retrieved only clade B *amoA* gene sequences from the forest soil (Table 2). The widespread occurrence of comammox *Nitrospira* in the analyzed samples is consistent with the previously reported presence of mostly uncultured and uncharacterized *Nitrospira* members in the respective habitat types (e.g. Daims *et al.*, 2001; Martiny *et al.*, 2005; Ke *et al.*, 2013; Pester *et al.*, 2014). Future studies should determine which fraction of these environmental *Nitrospira* are strict NOB or comammox, respectively. The presence of multiple comammox OTUs, and in particular the co-occurrence of both clade A and clade B comammox, in several samples (Table 2) indicates ecological niche partitioning that enables the coexistence of different comammox strains. For co-occurring *Nitrospira* in WWTPs, niche partitioning based on the preferred nitrite concentrations, on the capability to utilize formate as an alternative substrate, and on the tendency to co-aggregate with AOB has already been demonstrated (Maixner *et al.*, 2006; Gruber-Dorninger *et al.*, 2015). It is tempting to speculate that underlying mechanisms of niche differentiation of comammox *Nitrospira* could also be different substrate (ammonia) concentration optima and alternative energy metabolisms, such as the oxidation of formate and of hydrogen (Koch *et al.*, 2014; Koch *et al.*, 2015).

### 3.3 Quantification of comammox *amoA* genes by quantitative PCR

Since the discovery of AOA (Könneke *et al.*, 2005), numerous studies compared the abundances of AOB and AOA, and tried to assess the contributions of different groups of ammonia oxidizers to nitrification in various engineered and natural environments (e.g. Leininger *et al.*, 2006; Chen *et al.*, 2008; Jia and Conrad, 2009; Mussmann *et al.*, 2011; Daebeler *et al.*, 2012; Ke *et al.*, 2013; Bollmann *et al.*, 2014). To adequately investigate the niche differentiation and labor partitioning between ammonia oxidizers, it will be necessary to include comammox *Nitrospira* in such comparisons. This will require a robust method for the rapid and accurate quantification of comammox *Nitrospira.* Thus, we established qPCR assays using the equimolar primer mixtures that target comammox *amoA* clade A or clade B, respectively. Both assays had a high efficiency, accuracy and sensitivity, as the quantification of as few as ten copies of comammox *amoA* standards was achieved (Fig. S1).

As proof of applicability, we quantified comammox *amoA* clade A and clade B genes, alongside with archaeal and bacterial *amoA* genes, in five different sample types: one activated sludge sample (WWTP VetMed, Vienna); pasty iron sludge from the riser pipe of GWW Wolfenbüttel, Germany; a trickling filter sample from the DWTP Friedrichshof, Germany; paddy rice soil from Vercelli, Italy and a beech forest soil from Klausen Leopoldsdorf, Austria (Fig. 2, Table S2). Melting curve analyses confirmed the specificity of both newly established comammox *amoA* targeted assays (Fig. S2). Amplification of nontarget DNA occurred only in those samples that lacked the respective target organisms of the primers according to the aforementioned results of the end point PCR (i.e., WWTP VetMed for clade B and Klausen-Leopoldsdorf forest soil for clade A comammox) and was easily detectable based on a distinct slope of the melting curves of the unspecific amplicons (Fig. S2). The copy numbers of clade A and clade B comammox *amoA* genes were in the same order of magnitude in the samples containing both groups (Fig. 2, Table S2). Interestingly, the predominance of the different groups of ammonia oxidizers varied strongly among the analyzed samples (Fig. 2). In the activated sludge from WWTP VetMed, the gene abundances of betaproteobacterial and comammox clade A *amoA* were significantly different (p<0.01). Comammox clade A *amoA* copy numbers accounted for 14 to 34% of the estimated total *amoA* copy number (Table S2), indicating that comammox *Nitrospira* could be functionally relevant for nitrification in this system. The *amoA* qPCR-based detection of both comammox *Nitrospira* and AOB, as well as the absence of AOA, in WWTP VetMed are consistent with an earlier analysis of the same sample based on metagenomics and rRNA-targeted FISH (Daims *et al.*, 2015).The qPCR-based dominance of clade A and B comammox *Nitrospira* over AOB (p<0.01) in GWW Wolfenbüttel (Fig. 2, Table S2) is in line with the previous metagenomic detection of both clades in samples from the same system (Daims *et al.*, 2015). Altogether, the results obtained in this and other studies (Daims *et al.*, 2015, van Kessel *et al.*,2015; Palomo *et al.*, 2016; Pinto *et al.*, 2016; Bartelme *et al.*, 2017; Wang *et al.*, 2017) for wastewater and drinking water treatment plants suggest that comammox *Nitrospira* have to be considered in future studies of nitrification in engineered systems. Likewise, the relative abundances of comammox *amoA* genes in comparison to those of AOA and AOB in the two soils (Fig. 2) indicate that comammox organisms should not be neglected in research on terrestrial nitrifiers.

**Figure. 2.**
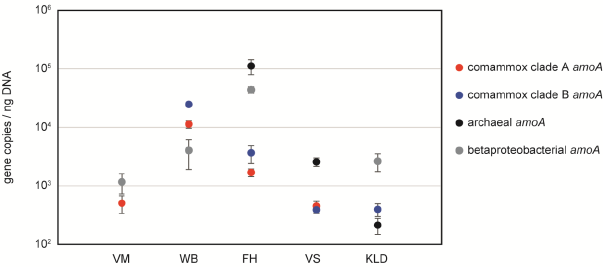
Abundances of *amoA* genes from comammox *Nitrospira*, AOA and AOB in selected samples from various environments. VM – WWTP VetMed; WB – GWW Wolfenüttel; FH – DWTP Friedrichshof; VS – Vercelli rice paddy soil; KLD – Klausen-Leopoldsdorf beech forest soil.

## 4 Summary and Outlook

The newly developed primer sets were designed for maximal coverage of the known comammox *Nitrospira* lineages, high specificity for comammox *amoA* genes, and a broad range of applications including high-throughput amplicon sequencing and qPCR. By using these primer sets, we detected the presence of comammox *Nitrospira* in a variety of habitats and found their *amoA* genes to be of comparable abundance as the *amoA* of other ammonia oxidizers. In this methodological study, our primary goal was to evaluate the applicability of the newly designed primers for end point PCR and qPCR with a selection of different sample types. A much broader census of comammox *Nitrospira* in the environment is needed before general conclusions on their distribution and importance for nitrification can be drawn. However, as was previously shown for AOA in a WWTP (Mussmann *et al.*, 2011), high *amoA* gene copy numbers do not always indicate that the respective population is functionally important for ammonia oxidation. Aside from the possibility that abundant putative nitrifiers grow on alternative substrates, one can also not exclude *amoA* gene quantification biases caused by extracellular DNA, which may for example be secreted to a different extent by the phylogenetically diverse nitrifiers. Moreover, at least in AOA a basal *amoA* transcription can occur even in the absence of detectable ammonia-oxidizing activity (Mussmann *et al.*, 2011; Ke *et al.*, 2013). In contrast, pronounced shifts in the *amoA* transcription levels upon certain environmental stimuli, such as a changing availability of ammonium, likely are useful activity indicators for AOA, AOB, and comammox *Nitrospira.* Although not tested yet in this context, the new comammox amoA-specific primers should in principle be suitable for reverse transcription-qPCR of *amoA* and should thus allow the quantification of comammox *amoA* transcripts in environmental samples and during incubation experiments. Thus, the primers can serve as a tool for a broad range of study designs to elucidate the ecological significance of comammox *Nitrospira* in comparison to other ammonia oxidizers.

## Author contributions

P.P., S.L., and H.D. conceived the research idea. P.P., S.L., H.D., M.A.H.J.v.K, M.S.M.J, and M.W. contributed to the development of the research plan and project goals. C.W.H. performed database mining, and C.W.H. and P.P. compiled the pmoA/amoA database used for primer design by P.P. C.S., L.P., P.P., A.D., and M.S. performed laboratory work, and contributed to data analysis. S.L., H.D., M.A.H.J.v.K, M.S.M.J, and M.W. contributed sources of project funding. P.P. and H.D. were the primary authors writing the manuscript, while all other co-authors were involved in writing and editing of the manuscript.

## Acknowledgments

The authors would like to thank Bernd Bendinger, Kay Bouts, Stephanie Eichorst, Mohammad Ghashghavi, Hannes Schmidt, Jasmin Schwarz, Annika Vaksmaa, Weren de Vet, Julia Vierheilig, Dagmar Woebken, and Thomas Zechmeister for providing sample material. Ping Han, Bela Hausmann, and Roey Angel are acknowledged for valuable discussion and advice with the qPCR assay setup. Andreas Pommerening-Röser is acknowledged for providing the *N. nitrosa* Nm90 isolate.

